# Visual areas exert feedforward and feedback influences through distinct frequency channels

**DOI:** 10.1101/004804

**Authors:** Andre M. Bastos, Julien Vezoli, Conrado A. Bosman, Jan-Mathijs Schoffelen, Robert Oostenveld, Jarrod R. Dowdall, Peter De Weerd, Henry Kennedy, Pascal Fries

## Abstract

Visual cortical areas are thought to form a hierarchy and to subserve cognitive functions by interacting in both feedforward and feedback directions^1^. While feedforward influences convey sensory signals, feedback influences modulate brain responses to a given sensory stimulus according to the current behavioural context. Many studies have demonstrated effects of feedback influences on feedforward driven responses^2^ and on behaviour^3^. Also, anatomical projections in both directions have been identified^1, 4^. However, although these studies have revealed the anatomical paths and the neurophysiological consequences of influences in both directions, the neurophysiological mechanisms through which these influences are exerted remain largely elusive. Here we show that in the primate visual system, feedforward influences are carried by theta-band (∼4 Hz) and gamma-band (∼60-80 Hz) synchronization, and feedback influences by beta-band (∼14-18 Hz) synchronization. These frequency-specific asymmetries in directed influences were revealed by simultaneous local field potential recordings from eight visual areas and an analysis of Granger-causal influences across all 28 pairs of areas. The asymmetries in directed influences correlated directly with asymmetries in anatomy and enabled us to build a visual cortical hierarchy from the influence asymmetries alone. Across different task periods, most areas stayed at their hierarchical position, whereas particularly frontal areas moved dynamically. Our results demonstrate that feedforward and feedback signalling use different frequency channels, which might subserve their differential communication requirements and lead to differential local consequences. The possibility to infer hierarchical relationships through functional data alone might make it possible to derive a cortical hierarchy in the living human brain.

---

Many aspects of cognitive performance can only be explained through the concept of feedback influences. For example, reaction times are shortened when stimulus locations are pre-cued and attention can be pre-directed, an effect that cannot be explained if only constant feedforward input is considered^3^. Numerous neurophysiological studies have demonstrated the effects of feedback influences on neuronal activity^2^, yet the mechanisms through which feedback influences are exerted remain elusive. Anatomical studies have revealed that structural connections in the feedforward direction, i.e. from the primary sensory areas to higher order areas, are complemented by connections in the feedback direction^1, 4^. In addition, it is well established that feedforward and feedback connections follow a characteristic pattern with regard to cortical layers: Feedforward connections target the granular layer^1^; they originate preferentially in supragranular layers, and this preference is stronger for projections traversing more hierarchical levels, i.e. it is quantitatively related to the hierarchical distance^4^. Feedback connections avoid targeting the granular layer^1^; they originate preferentially in the infragranular layers, and again, this preference is stronger for projections traversing more hierarchical levels and is thereby quantitatively related to hierarchical distance^4^. These asymmetries have been used to arrange the visual cortical areas into a hierarchy^1, 4^, which has influenced many theories of cognition and brain function^5, 6^.

Recent studies have documented a neurophysiological asymmetry between cortical layers in visual cortex: While supragranular layers show local gamma-band synchronization, infragranular layers show local alpha/beta-band synchronization^7–9^. Local rhythmic synchronization can lead to inter-areal synchronization^10–12^, which has been proposed as a mechanism of effective inter-areal interaction^11, 13, 14^. Given that supragranular layers primarily send feedforward projections and infragranular layers primarily feedback projections, this leads to the hypothesis that inter-areal synchronization in the gamma-frequency band might mediate feedforward influences, and inter-areal synchronization in the beta-frequency band might mediate feedback influences.

To test this prediction, we recorded local field potentials (LFPs) from electrocorticography (ECoG) grids implanted onto the left hemispheres of two macaque monkeys (Fig. 1a,b and Extended Data Fig. 1) performing a visuospatial attention task (Extended Data Fig. 2 and Methods). The ECoG grid covered eight visual areas: V1, V2, V4, TEO, DP, 7A, 8L and 8M (lateral and medial parts of area 8/FEF)^11, 15, 16^. The 252 electrodes were assigned to cortical areas by co-registering intraoperative photographs with several macaque brain atlases^17^ (Extended Data Fig. 1). For the frequency bands analysed here, ECoG signals reflect neuronal activity from both superficial and deep cortical layers^18^. Signals from immediately neighbouring electrodes were subtracted from each other to construct local bipolar derivations (Extended Data Fig. 1), which we will refer to as “sites”. Between pairs of sites from different areas, inter-areal synchronization is quantified by the coherence metric (see Methods). For an example pair of areas, V1 and DP, the inter-areal coherence during visual stimulation and task performance (“post-cue” period, see Methods) revealed distinct peaks in the theta-, beta- and gamma-frequency bands (Fig. 1c). This spectral pattern was consistent across inter-areal site pairs (Extended Data Fig. 3; see Supplementary Discussion for an account of the absolute coherence values; Extended Data Figure 3 displays the theta peak particularly clearly, because it is based on 1 s data epochs, giving a 1 Hz resolution. For most analyses, we used 0.5 s epochs for consistency across task epochs and optimal data use, and the resulting 2 Hz resolution flattens out the theta peak). We determined frequency-specific directed influences by calculating Granger-causal (GC) influences between all possible inter-areal pairs of sites^19^. The spectrum of GC influences of site 1 onto site 2 quantifies, per frequency, the variance in site 2 that is not explained by the past of site 2, but by the past of site 1. For our example pair of areas, the influence of V1 onto DP is a feedforward influence^4^ and it exceeded the feedback influence in the theta and gamma bands, while the feedback predominated in the beta band (Fig. 1c).

**Figure 1.**
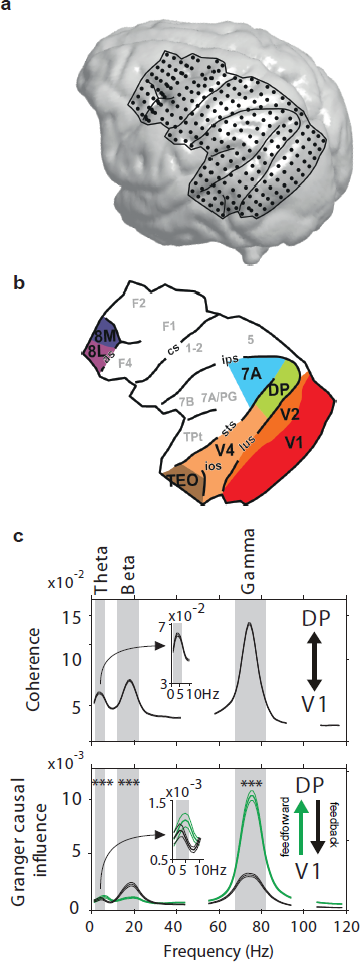
ECoG electrode distribution and interaction spectra for an example pair of areas. **a,** Rendering of the brain of monkey 1 based on structural MRI scans. Lines indicate the boundary of the covered brain region and the major sulci, and dots indicate the 252 subdural electrodes. **b,** Parcellation of ECoG-covered regions into cortical areas. **c,** Coherence (upper panel) and GC influence (lower panel) spectra for an example pairs of areas: V1 and DP. Values in the range from 45-55 Hz and 95-105 Hz are masked because of residual line noise.

**Figure 2.**
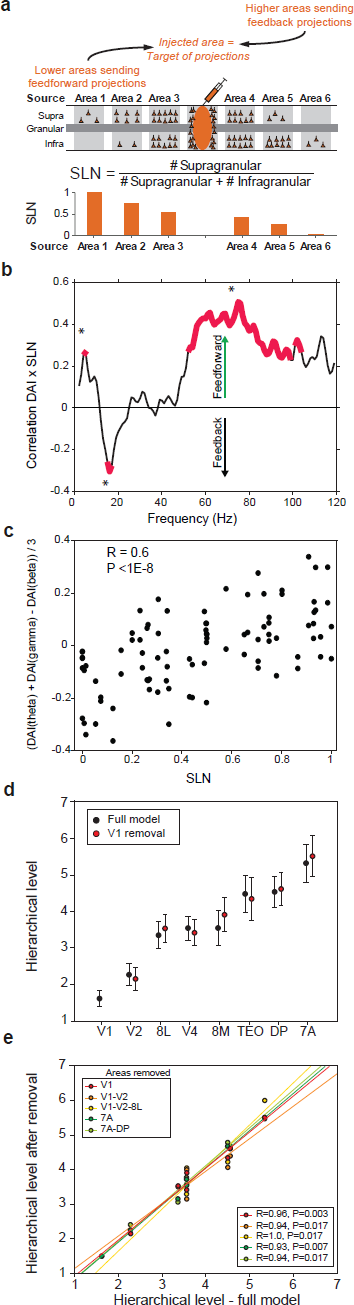
Granger-causal influences correlate directly with anatomy and establish a hierarchy. **a,** Schematic of retrograde anatomical tracing method and calculation of SLN values. Retrograde tracer is injected into a target area and labels neurons in several source areas projecting to the target area. Source areas hierarchically lower (higher) than the target area have a progressively higher (lower) proportion of labelled neurons in the supragranular layers, i.e. the lower (higher) the area, the higher (lower) the SLN value. **b**, Correlation between SLN and DAI per frequency (across area pairs and monkeys) revealed the main direction of frequency-specific influence. Three frequency-bands were significantly correlated (randomization test with multiple comparison correction across frequencies): Theta (4-5 Hz), Beta (15-17 Hz) and Gamma (∼50-100 Hz); theta and gamma influences predominated in the feedforward direction and beta influences in the feedback direction. To optimally capture the theta peak, this analysis was performed on 1 s data epochs. **c,** Correlation between SLN and the DAI combined across theta, beta and gamma bands as specified on the y-axis. Only SLN values based on at least 10 labelled neurons were included. **d,** Black dots: Hierarchical levels for all areas, derived by taking each area in turn as target and assigning the hierarchical level to the other areas based on their GC influences to the target. Error bars show the SEM across target areas. Red dots: Hierarchical levels after removing V1, revealing immunity to this manipulation. **e,** Red dots: Hierarchical levels of the full model versus one with V1 removed. Other colours: Corresponding analyses after removing more areas from the lower or upper end of the hierarchy.

**Figure 3.**
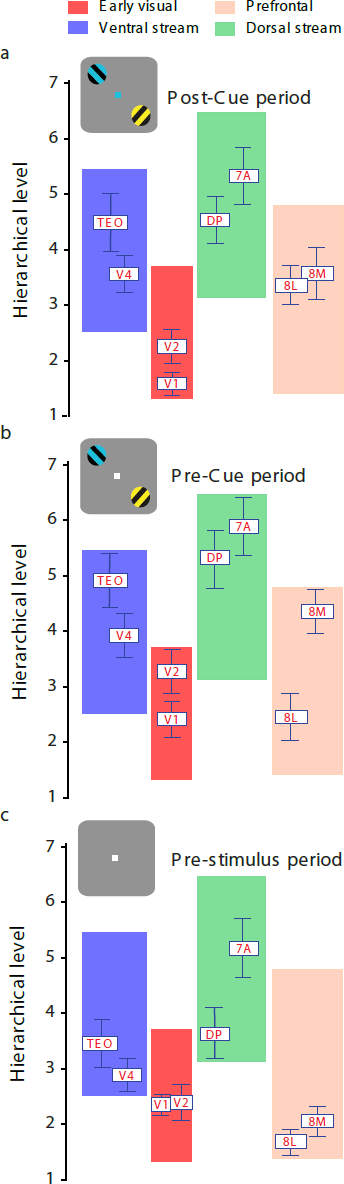
The functional hierarchy is dynamic. The dynamics of the functional hierarchy with cognitive context is shown through three main periods of the task. (**a**) The post-cue period, when the stimulus was on and the attentional cue had been given and attention had been deployed. (**b**) The pre-cue period, when the stimulus was on, but the attentional cue had not yet been given. (**c**) The pre-stimulus period, when the animal was fixating, but the stimulus was not yet presented. Each area's mean hierarchical position is depicted relative to the others. Error bars indicate standard error of the mean in the hierarchical position across the different areas taken as targets.

To test whether this pattern held generally, we related GC influences to anatomical connections, specifically to a metric of their feedforward/feedback directedness. When retrograde tracer is injected into a target area, the target-projecting neurons are labelled in different source areas. If a source area is providing feedforward (feedback) input to the target area, the proportion of **<u>s</u>**upragranular **l**abelled **<u>n</u>**eurons (relative to supragranular plus infragranular labelled neurons) is high (low)^4^. This SLN metric quantifies the degree to which an inter-areal anatomical connection is feedforward or feedback (Fig. 2a). We aimed at directly relating the SLN to a graded metric of functional asymmetry. We used the Granger-causal inter-areal influences to define a **<u>d</u>**irected-influence **<u>a</u>**symmetry **<u>i</u>**ndex (DAI). Similar to the anatomical approach, we considered each area consecutively as the target of directed influences. With regard to this target, the influence from any given source area to the target was considered the “inflow”, and the influence of the target onto the source area the “outflow”. The DAI was then defined as (outflow-inflow)/(outflow + inflow). Across all target-source area pairs, we correlated the DAI with the corresponding SLN values (using DAI values from two monkeys with ECoG recordings and SLN values from an independent set of 25 monkeys). Because the DAI is defined per frequency, the DAI-SLN correlation was also determined per frequency and the resulting correlation spectrum is shown in Figure 2b. This spectrum shows significant positive DAI-SLN correlations for theta and gamma, and a significant negative correlation for beta. A positive DAI-SLN correlation shows that increasing SLN, i.e. decreasing hierarchical level of the source area, corresponds to increasing DAI, i.e. relatively more outflow. Therefore, a positive (negative) DAI-SLN correlation for a given frequency indicates that this frequency channel conveys feedforward (feedback) influences. Thus, the correlation spectrum demonstrates that feedforward influences are conveyed through theta- and gamma-frequency channels, and feedback influences are conveyed through a beta-frequency channel. Similar results were obtained for the other task periods (Extended Data Fig. 4).

**Figure 4.**
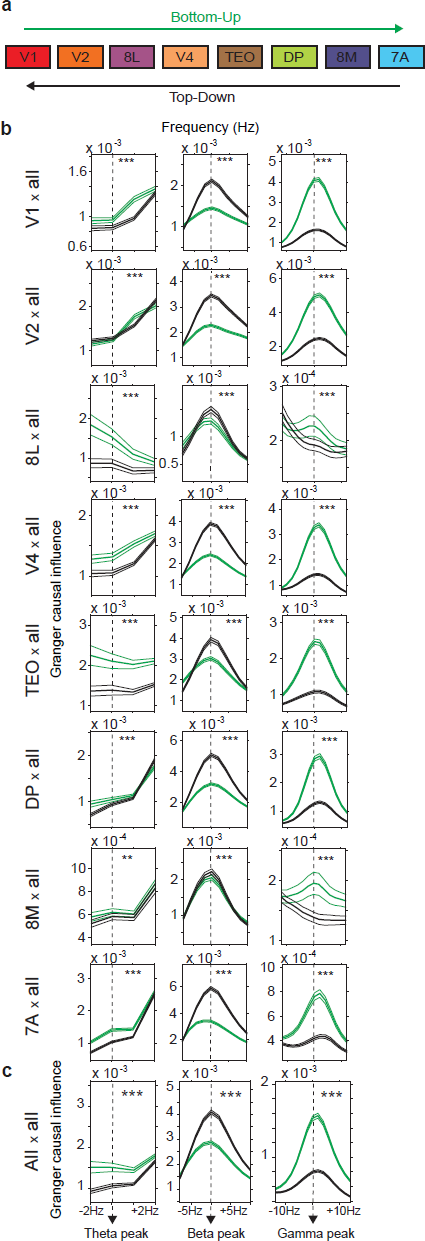
Granger-causal influence asymmetries. **a,** Hierarchical ranking of the recorded visual areas according to the most recent anatomical hierarchical model^4^. **b,** Each area was taken as target area, as indicated to the left of the panels, and the GC influences between that area's sites and all other sites were averaged. GC influence spectra were sorted into bottom-up (green) and top-down (black) directions according to the anatomical hierarchical ordering shown in **a**. Spectra were averaged across monkeys after aligning frequency peaks. **c,** Same as **b**, but grand averaging across all target areas.

The pattern of anatomical feedforward and feedback connections across all pairs of visual areas is largely consistent with a global hierarchy, in which each area occupies a hierarchical level, which in turn defines a given inter-areal influence as either bottom-up or top-down. Please note that such a hierarchy is a global model fitted to the area-pair-wise relations, and the hierarchy-derived bottom-up (top-down) relationships agree only partly with the inter-areal feedforward (feedback) relationships. The correlations between the anatomical SLN metric and the functional DAI metric suggest that it might be possible to construct a hierarchy of visual cortical areas from DAI values alone, which would demonstrate that not only the anatomical, but also the functional relations across many pairs of areas are consistent with a global hierarchy. To explore this, we first used the post-cue period and combined all evidence available in the DAIs across the frequency spectrum, by averaging the DAIs of the theta-, beta- and gamma-frequency bands, after inverting the sign of the beta-band DAI, because of its negative correlation to SLN. This multi-frequency-band DAI was strongly correlated with the SLN across all pairs of areas (R = 0.6, P < 1e^−8^, using Spearman rank correlation here and in the following correlation tests) (Fig. 2c). We proceeded to construct the functional hierarchy as follows: The multi-frequency-band DAI values were (1) normalized to range from zero to ten, (2) shifted for each target area consecutively such that the smallest value for the lowest hierarchical level was one, and (3) averaged across all target areas. Figure 2d (black dots) shows for the eight areas the resulting hierarchical levels and their standard error across area-pairs and monkeys. To further probe the robustness of the functional hierarchy, we removed one or multiple areas and built the functional hierarchy on the remaining areas. The red dots in figure 2d show that removal of V1 leaves the hierarchical positions of the remaining 7 areas essentially unchanged. These positions were plotted against the positions from the full model as red dots in figure 2e, demonstrating a strong correlation (R = 0.96, P = 0.003). This correlation remained significant even when we removed up to three areas from the lower end of the hierarchy, or up to two areas from the upper end (Fig. 2e, other colours).

The complete functional hierarchy is shown again in Figure 3a and strongly correlates with the most recent anatomical hierarchy of visual cortex^4^ (R = 0.93, P = 0.002). The functional hierarchy is defined by Granger-causal influences, with the intriguing consequence that it might change dynamically. This would require dynamic changes in GC influences between areas, which have been described e.g. between FEF and V4 during the course of task performance^10^. Therefore, we investigated, whether the functional hierarchy changed across different task periods. We found that the post-cue hierarchy was already largely present during the pre-cue period (Fig. 3b). Areas V1, V2, V4, TEO, DP and 7A were arranged in their well-established order. Yet, 8L, the lateral part of FEF, assumed a lower level in the pre-cue period (Fig. 3b). In the pre-stimulus period (Fig. 3c), 8L moved to the bottom of the hierarchy and 8M joined in the immediate neighbourhood. Furthermore, V1, V2 and V4 moved closer together. These analyses demonstrate that the DAI-based functional hierarchy is not fixed like anatomy-based hierarchies. The most recent anatomy-based hierarchy showed an R = 0.93 correlation to the post-cue functional hierarchy (P = 0.002), an R = 0.91 correlation to the pre-cue functional hierarchy (P = 0.005), and no significant correlation to the pre-stimulus functional hierarchy (P = 0.2). Once the stimulus is present, inter-areal influences are most likely exerted in both bottom-up and top-down directions. Anatomical connections in the two directions are present at all times. This might explain why the anatomical hierarchy correlates particularly well with the functional hierarchy during stimulation.

Thus, like in anatomy, the pattern of feedforward and feedback GC influences across all pairs of simultaneously recorded visual areas is largely consistent with a global hierarchy. Also as in anatomy, the global hierarchy agrees only partly with the area-pair-wise GC influence pattern (Extended Data Fig. 5–7). Yet, when separate tests (Bonferroni corrected across all tests) were performed per area pair, frequency band and monkey, significant differences between GC influences in the two directions agreed with the anatomical hierarchy in the majority of cases (47 of 61, or 77%, P < 0.001 across all tests; P < 0.02 for theta, P < 0.03 for beta, P < 0.005 for gamma; binomial tests). When we defined each area in turn as the target area and averaged its GC influences to all other areas, theta-band influences were more bottom-up directed for 7 of 8 target areas, beta-band influences were more top-down directed for all target areas, and gamma-band influences were more bottom-up directed for all target areas (Fig. 4a,b). In the grand average across all 28 pairs of areas and both animals this pattern was highly significant (Fig. 4c, P = 0 for each of the three frequency bands). Additional analyses showed that this pattern was not due to observation noise^20^ (Extended Data Fig. 8a,b) or the bipolar derivation scheme (Extended Data Fig. 8c,d). Also, conditional GC influence analysis^21^ left the pattern of results unchanged for gamma and beta, and suggested the involvement of larger networks for theta (Extended Data Fig. 9).

Finally, we tested the prediction that top-down beta-band influences are enhanced when a cognitive task requires stronger top-down control. Top-down control is expected to be enhanced by selective attention. Indeed, when selective attention was directed to the contralateral as compared to the ipsilateral stimulus, top-down beta-band GC influences were enhanced in the grand average (P < 0.001) and in all pairs of areas with a significant attention effect (N = 13, P < 0.0005, binomial test).

In summary, we have shown that among primate visual cortical areas, feedforward communication utilizes the theta and gamma band, and feedback communication the beta band. As gamma-band (beta-band) synchronization predominates in superficial (deep) cortical layers^7–9^, these asymmetries in directed influences are likely related to the laminar pattern of inter-areal anatomical connections. Future studies might test this directly with simultaneous multi-area multi-layer recordings of LFP and spikes, and might extend the coverage to more cortical and subcortical structures, and the previous laminar analyses^7–9^ to the theta band. Regarding the theta band, we note that the visual cortical theta rhythm is partly locked to microsaccades^22^. Therefore, theta-rhythmic microsaccades with corresponding retinal image motion and subsequent visual responses might contribute to the feedforward GC influences in the theta band. For the gamma band, an analysis that excluded microsaccade effects left the pattern of GC influences unchanged (Extended Data Fig. 10).

It is likely that feedforward and feedback inter-areal influences need to fulfil different requirements, which might be met by synchronization in different frequency bands with their different time scales. For example, it is conceivable that inter-areal synchronization entails higher energetic costs for gamma than beta^23^, and bottom-up signalling might be equipped with gamma-band activity to achieve higher communication throughput. At its target structure, an input might have differential effects solely due to the rhythm through which it has been transferred. For example, target cells and/or local circuits with resonant properties in particular frequency bands might be addressed differentially by inputs with different rhythms^24–27^. In that sense, the frequency band through which an input is mediated might functionally tag that input to trigger differential further processing.

## Methods Summary

Two adult male rhesus monkeys performed a visual attention task, during which they fixated a central spot and released a bar when the behaviourally relevant stimulus underwent a shape change (Extended Data Fig. 2). Behavioural relevance was assigned on a trial-by-trial basis with a centrally presented cue. Two stimuli were presented, one in the lower right visual hemifield, and one in the upper left visual hemifield. Neuronal signals were recorded from the left hemisphere in two monkeys using subdural ECoG grids consisting of 252 electrodes (1 mm diameter), which were spaced 2-3 mm apart^11, 15, 16^. Data were recorded in 9 sessions in monkey 1 and 14 sessions in monkey 2. The post-cue analysis used the time period from 0.3 s after cue onset until the first shape change in one of the stimuli. Only trials with a correct behavioural report were used. For each trial, this period was cut into non-overlapping 0.5 s data epochs. This resulted in 3874 epochs for monkey 1 and 3492 epochs for monkey 2. For both the pre-stimulus and pre-cue periods, there were 4239 and 4396 epochs of 0.5 s in monkey 1 and 2, respectively. For each site and recording session, the data epochs were normalized by their standard deviation and subsequently pooled across sessions. Data epochs were multitapered using three Slepian tapers and Fourier-transformed^28^. The epoch lengths of 0.5 s resulted in a spectral resolution of 2 Hz, the multitapering in a spectral smoothing of ± 3 Hz. Where mentioned explicitly, we used Hann-tapered 1 s epochs for 1 Hz spectral resolution. The Fourier transforms were the basis for calculating the coherence spectra and for calculating the GC influence spectra through non-parametric spectral matrix factorization^19^. The non-parametric estimation of GC influences spectra has certain advantages over parametric approaches, e.g. it does not require the specification of a particular model order.

## Acknowledgments

This work was supported by Human Frontier Science Program Organization Grant RGP0070/2003 (P.F.), Volkswagen Foundation Grant I/79876 (P.F.), the European Science Foundation European Young Investigator Award Program (P.F.), the European Union (HEALTH-F2-2008 - 200728 to P.F.), the LOEWE program (“Neuronale Koordination Forschungsschwerpunkt Frankfurt” to P.F.), the Smart Mix Programme of the Netherlands Ministry of Economic Affairs and the Netherlands Ministry of Education, Culture and Science (BrainGain to P.F., R.O. and J.M.S.), The Netherlands Organization for Scientific Research Grant 452-03-344 (P.F.), the National Science Foundation Graduate Student Fellowship 2009090358(A.M.B.), a Fulbright grant from the U.S. Department of State (A.M.B.), ANR-11-BSV4-501 (H.K.), and LabEx CORTEX (ANR-11-LABX-0042 to H.K.). A.M.B. would like to thank G.R. Mangun and W.M. Usrey for support.

## Author Contributions

C.A.B. and P.F. designed the experiments; C.A.B. trained the monkeys and recorded the electrophysiological data; P.F., P.D.W. and C.A.B. implanted the monkeys; J.M.S., R.O., C.A.B. and A.M.B. wrote analysis programs; A.M.B., J.V., and J.M.S. performed analyses with the help of R.O. and with the advice of P.F.; H.K. provided the anatomical data (SLN); J.R.D. contributed to the microsaccade analysis; A.M.B., J.V., and P.F. wrote the paper in collaboration with P.D.W, C.A.B., J.M.S., and H.K.

## Author Information

Updates, atlases and additional information concerning the anatomical dataset that was used for this work is available at www.core-nets.org. Reprints and permissions information is available at www.nature.com/reprints. The authors declare no competing financial interests. Correspondence and requests for materials should be addressed to P.F., A.M.B., J.V., or C.A.B.

**Supplementary Information** is linked to the online version of the paper at www.nature.com/nature.

